# Isolation and Identification of possible pathogenic bacteria on carrots sold at community market, Ahmadu Bello University, Zaria

**DOI:** 10.1101/2023.09.29.559988

**Authors:** Blessing Ojodomo, Saaondo James Ashar

## Abstract

Carrots (*Daucus carota*) are root vegetables that can be purple, red, yellow or white colour based on their pigmentation and variety. Due to the awareness of its high nutritional content such as vitamins, minerals and carotene, they are mainly consumed raw, possibly exposing one to the risk of being infected. Studies on the pathogenic Bacteria associated with fresh carrots sold in the community market at Ahmadu Bello University, main campus Samaru Zaria, were carried out using cultural techniques. Salmonella Shigella agar, MacConkey agar and Eosin methylene blue agar were the growth media for the isolation of the pathogenic Bacteria. Twelve fresh carrots were randomly bought from different food vendors in the community market, and then 10g of the outer layer (epidermal) scrapings were collected from each sample and homogenized in 90 ml of buffered peptone water. It was then serially diluted and 0.1ml of each 10^4^ and 10^5^ dilutions were plated on EMB, SSA, and MacConkey agar. Isolates were identified by gram staining and biochemical tests. The bacteria were identified to be *Proteus* species, *Citrobacter* species, *Klebsiella* species, and *Salmonella* species. *Proteus* spp was predominantly isolated among the bacterial isolates (41%) followed by *Citrobacter* spp (25%), *Klebsiella* spp (17%) and *Salmonella* spp (17%). These organisms may have been introduced to the carrots during growth, harvesting, handling, storage and distribution. The presence of the organisms is a public health risk because of the diseases known to be caused by them. It is therefore imperative that adequate hygienic practices be put in place during the cultivating and handling of carrots. A high level of susceptibility was observed in Ciprofloxacin at (100%) and Augmentin at (50%), while the high level of resistance was observed to be high in Gentamicin at (100%).

## CHAPTER ONE

### INTRODUCTION

#### 1.1 Background to the study

Carrots (Daucus carota) is a biennial herbaceous species, it is part of the Apiaceae Family. Carrots are classified into two mainly Western carrots and Eastern carrots and this is based on carrot pigmentation. The origin of western carrots is not yet known while the eastern carrots is said to originate from Afghanistan. Most carrots root is purple and some are yellow. The leaves are slightly dissected and roots branched. Currently the more widely cultivated carrots in the world are the orange carrots and are more popular. (Que, F., Hou, Xl, Wang, Gl *et al*., 2019)

Carrots are grown in sandy loam or silt loam soil most at times to enhance water holding capacity and drainage. Planting carrots in raised beds can further help in proper water drainage. Carrots need soil that has adequate air and water drainage because wet and compacted soils can cause a deformed growth. The temperature of the soil three inches below the surface should be 50°F or lower. Carrots can withstand PH ranging from 5.5 to 8.0 because there are hard crops, however, light sandy soil with a neutral PH and under full sun exposure, this is opposite to very clay-like or wet, chalky soil. Tillage of soil is done to loosen the compacted ground before seeding. To have the best root development and growth, carrots should have approximately 18-24 inches of ell-tilled soil that has adequate drainage. Abnormal shaped or forked carrots that are unmarketable are grown due to the presence of pebbles and stones in the soil. Pythium root dieback, nematodes, and exposure to frost are other factors that could cause stubbed of forked roots (Anupama *et al*., 2020)

Carrots are crop that are able to adequately extract nutrient from the soil due to their deep-rooting nature. It is necessary for soil test to be carried out before planting and throughout development to measure soil nutrient such as Nitrogen, Potassium, Phosphorus, Magnesium, Manganese, Boron and Sulphur. However, nutrient can be added before seeding and during crop maturation with the use of side dressing or broadcasting. Precaution should be taken as excess nitrogen in the soil causes root cracking during harvest. Due concern for food safety and high nitrogen, addition of fresh manure is not advisable. (Pensack-Rinehart and Buning 2015).

Carrots are the most important crop in the Apiaceae family. Carrots was first used for medicinal purpose and later used as food. Orange carrots the most popular was cultivated in 15^th^ and 16^th^ centuries in central Europe. The reason for popularity of orange carrots was because it was observed to contain high ProVitamine A. The major Antitoxidant found in carrots are Carotenoids and Anthocyanin. Yellow carrots are highly rich in Alpha and Beta carotene and rich in ProVitamine A (da Silva Dias 2014). Lutein present in carrots is responsible for its yellow color and plays an important role in macular degeneration prevention. Carotene level gradually increases with growth and is more concentrated at the corticle than the core. Carrots have high nutritional value. It is a good source of dietary fiber and of trace minerals molybdenum (Nicolle *et al*., 2004).

Carrots is a root vegetable that contain carotenoid, flavonoids, polyacetylenes, vitamins and minerals, all of these possess numerous nutrition and health benefits. They were an old adage that carrots are good for the eyes. Carotenoid, polyphenol and vitamins present in carrots act as antitoxidant, anticarcinogenics, and immunoenhancers. Antidiabetic cholesterol and cardiovascular disease, lowering, antihypertension, hepatoprotective, renoprotective and wound healing benefits of carrots also have been reported (da Silva Dias 2014).

Processed vegetables, the spoilage of horticultural products justifies the use of preservative techniques. This processing not only adds value to the products, but as well makes the products more convenient to be consumed by consumers. Consumers requested for high quality, a fresh, nutritive and conveniently prepared vegetable has increased so much in the recent years. This has led to the development of lightly processed vegetables. Preparation of lightly processed carrots is done by peeling the epidermal layer of the carrot roots; this is one of the most popular products that are available in the United States. One of the disadvantages of this processing method is that it makes carrots susceptible to different physiological changes that cut short their shelf-life. The peeling of the epidermal layer of the carrots increase these potential for carotene oxidation during storage, this also may further increase the respiration of carrots tissue resulting in increased degradation of protein, carbohydrates, lipids and the development of off-flavors (Peiyin and Barth 1998).A new protective layer called white blush is developed when the epidermal layer is peeled off and this result in dehydration and lignification on the carrots surface (Bolin and Huxsoll, 1991).

#### 1.2 Statement of Research Problem

Increase in awareness of the health benefits of vegetables has resulted in an increase in consumption. Many vegetables are consumed raw to retain the natural taste and heat labile nutrients. It is claimed that Microbes are found all over the globe with some few exceptions, including sterilized surfaces. They include normal flora that is nonpathogenic, which contribute to the larger percentage and pathogenic species which are few (Gadafi et al., 2020). The safety of raw vegetables is a great concern. This research and experiment are therefore centered on investigating and analyzing incidence of pathogenic bacteria on fresh carrot, to also know possible food borne bacteria pathogen on carrots (Anupama *et al*., 2020).

#### 1.3 Justification

Carrots are root vegetables that are highly consumed in every family. It is essential to health because of its high nutritional value. It provides nutrients such as vitamins and minerals and also is of medical important. Carrots are liable to contamination from various sources such as soil, man, water, air, and insects (Yong, 2014). Therefore, Isolation and identification of pathogenic bacteria from fresh carrots is necessary, to enlighten consumer of various ways of hygienic practices that leads to reduction of microbial load and a determination of the antibiotic susceptibility patterns of the isolates in case of food borne outbreak in the country (Anupama *et al*., 2020).

#### 1.4 Aim

The aim of this research was to isolate and identify possible pathogenic bacteria on carrots sold in community market Ahmadu Bello University Zaria.

#### 1.5 Objectives

The objectives of this research were:

i. to isolate and identify possible pathogenic bacteria on carrots sold in community market, Ahmadu Bello University, Zaria.
ii. to determine antimicrobial susceptibility patterns of the pathogens from carrots sold in community market, Ahmadu Bello University, Zaria

## CHAPTER TWO

### 2.0 LITERATURE REVIEW

#### 2.1 Carrot

##### 2.1.1 Origin and Domestication

The carrot (*Daucus carota)* is a root vegetable, usually orange in color, though purple, black, red, white, and yellow cultivars exist. By the existence of orange carrots, purple root color was apparently more common in eastern regions, yellow more common in the west. Eastern carrots tend to have less deeply divided leaflets with heavy leaf pubescence in some cultivars. For any carrot production, early flowering is unsatisfactory, eastern carrots have a greater tendency toward early flowering than western carrots, likely due to the somewhat warmer climates over the eastern production range. Beyond the yellow, purple, and orange root colors, eastern carrots have long included red-rooted types while western carrots included white-rooted types. Carrot use has also varied across production areas, with a more predominant use as animal forage in the east but largely human use as a root vegetable in the west (Philipp *et al.,* 2020).

Carrot is the most widely grown member of the Apiaceae or Umbelliferae. They are a domesticated form of the wild carrot, *Daucus carota* a native to Europe and Southwestern Asia. This diverse and complex plant family includes several other vegetables, such as parsnip, fennel, celery, root parsley, celeriac, arracacha, and many herbs and spices (Rubatzky *et al.,* 1999). The plant probably originated in Persia and was originally cultivated for its leaves and seeds (Wikipedia 2021). Underlying varietal distinctions based upon storage root color and shape is adaptation to cool versus warm growing temperatures. Carrot is categorized as a cool-season vegetable and the majority of effort on carrot breeding has been towards improving production in temperate regions where cool temperatures (< ∼10C) can stimulate early flowering or “bolting”. More recently there have been successful efforts in broadening the adaptation of carrot to warmer subtropical climates where excessive heat can retard plant growth, inhibit root color development, and stimulate the development of strong flavor in unadapted germplasm (Anupama *et al*., 2020). The ‘Brasília’ cultivar, for example, grows successfully in agricultural regions near the Equator. The development of temperate (late-flowering) and subtropical (early-flowering) types has resulted from a greater emphasis on ability to withstand early bolting in cooler climates for temperate types, in contrast to a greater emphasis on the ability to produce a marketable crop in warm climates for subtropical types (Philipp *et al.,* 2020). Subtropical carrots tend to grow faster than temperate types suggesting a complex interaction between root growth, flowering induction, and temperature that is not well understood. It should be noted that, unlike many crops, there is little evidence for a photoperiod effect on carrot root production and flowering so that the same cultivar theoretically could be grown anywhere in the world, if temperature requirements are met. In fact, many carrot cultivars are widely adapted and can be grown over such extreme production temperatures as represented by north of the Arctic Circle to highland subtropical climates. (Philipp *et al.,* 2020)

Like other plants of this family, carrot seeds are aromatic and consequently have long been used as a spice or herbal medicine. In fact, carrot seed was found in early human habitation sites as long as 3000 to 5000 years ago in Switzerland and Germany (Laufer, 1919). This seed is thought to be from wild carrot used for flavor or medicine. It also forms a major ingredient in the food processing industry, a significant constituent of cosmetic products and its image has long been used to symbolize healthy eating. The leaves are also consumed in salads and the seeds made into an herbal tea (John *et al.,* 2011).

In terms of both areas of production and market value, carrot is part of the top-ten most economically significant crops vegetable in the world (Rubatzky *et al*., 1999; Simon, 2000; Fontes and Vilela, 2003; Vilela, 2004). In 2005, world production approached 24 Mt on 1.1 million hectares. The total global market value of the more widely traded carrot seed crop has been estimated to be in the range of $100 million (Simon, 2000), but such estimates have little reliable data to confirm them and true value is likely much more. The development of cultivars adapted for cultivation in both summer and winter seasons on all continents has allowed a year-round availability of carrot products with relatively stable prices to consumers. Some production areas harvest crops year-round. Carrot improvement today includes several academic, private and government research programs around the world that work in concert with local, regional, and global industries. Both grower and consumer needs are addressed by public and private carrot breeders that incorporate modern technologies into the classical breeding process (Philipp *et al.,* 2020).

The genetic improvement of carrot has been an ongoing effort throughout its cultivation and domestication. Before the 20th century, carrot production was small scale in family or community gardens. A portion of the crop was likely protected in the field over winter with mulch, or the best roots saved in cellars were replanted the subsequent spring to produce a seed crop. There is no written record of what traits were evaluated or any other detail of the selection process in this period, but all domesticated carrot differs from its wild progenitors in forming larger, smoother storage roots, so it is clear that these traits also were improved through regular selection. Selection for low incidence of premature flowering was also necessarily among the most important traits selected during domestication, as it is now, since with the initiation of flowering, eating quality diminishes dramatically (Philipp *et al.,* 2020). One can say that color and flavor were primary selection criteria since they were the traits used to distinguish among carrots recorded by historians, cooks, and eventually seed catalogues. Carrot root color also changed dramatically during domestication. While wild carrot roots are white or very pale yellow, purple and yellow carrots were the colors of the first domesticated carrots. These were the only colors recorded until the 16th to 17th century when orange carrots were first described and soon came to be preferred in both the eastern and western production areas (Rubatzky et al. 1999, Simon, 2000). Banga compiled an extensive list of comments about carrots over history and while purple carrots were usually (but not always) regarded as better flavored than yellow, the dark stains they left on hands, cookware, and in cooking water raised negative comments by some authors. We do not know why early carrot breeders shifted their preference to orange types, but this preference has had a significant effect in providing a rich source of vitamin A, from alpha-and beta-carotene, to carrot consumers ever since. Soon after orange carrots became popular, the first named carrot cultivars came to be described in terms of shape, size, color, and flavor, and the first commercially sold carrot seed included reference to this growing list of distinguishing traits.

##### 2.1.2 Disease Resistance

Disease and pests limit carrot production to some extent in all carrot production regions. Leaf blights caused by *Alternaria dauci*, *Cercospora carotae*, and *Xanthomonas campestris* pv. *carotae*, powdery mildew (*Erisiphe heraclei*), carrot fly (*Psila rosae*), cavity spot (*Pythium* species and perhaps other pathogens), and several nematodes (e.g. *Meloidogyne* spp., *Heterodera carotae*, *Pratylenchus* spp.) are among the most widespread carrot diseases and pests, occurring worldwide. Several other pathogens and pests can cause very serious damage in more limited regions (Rubatzky *et al.,* 1999). Carrot breeders have relied upon natural infection in production areas where there is regular disease occurrence to make progress in selecting for genetic resistance for most diseases. Often highly susceptible cultivars or inbreeds are interspersed among entries to be tested in the field and in some cases natural inoculation is supplemented with inoculum from artificially infested plants. This approach has been used in selecting for resistance to *Alternaria* leaf blight (Boiteux *et al.,* 1993; Simon and Strandberg, 1998), and aster yellows (Gabelman *et al.,* 1994). For soil borne disease and pests, heavily infested disease evaluation plots have been established for *Meloidogyne incognita, M. javanica* (Vieira *et al.,* 2003), Methods for evaluating resistance to *Alternaria* leaf blight (Simon and Strandberg, 1998; Pawelec *et al*., 2006), cavity spot and *Rhizoctonia solani* resistance (Breton *et al.,* 2003), *M. hapla* (Wang and Goldman, 1996), and *M. javanica* (Simon *et al*., 2000) in controlled environments such as a greenhouse or growth chambers have also been developed.

##### 2.1.3 Consumer Quality

Selection for uniform orange color has been exercised by carrot breeders for the last century. The nutritional quality conferred by the provitamin. A carotenoid that account for the orange color of carrots has received the attention of carrot breeders since the 1960s beginning with extensive efforts of W.H. Gabelman and his students (Umiel *et al* 1972; Buishand *et al* 1979). As a result, selection has raised provitamin carotene content in typical U.S. carrot varieties by 70% between 1970 and 1992 (Simon, 1992). Yellow, purple, red, and white carrots have received a renewed level of interest in recent years as growers look for new niche markets and consumers become more aware of the nutritional benefits of pigments. To support selection with objective measurements of color, an evaluation tools have been developed (Surles *et al*., 2004).

Orange carrot color is primarily due to alpha-and beta-carotene, yellow and red carrot color are caused by carotenoids lutein and lycopene, respectively, and purple carrot color is caused by anthocyanins (Surles *et al.,* 2004). When no pigments accumulate, carrots are white. The commercial interest in carrots of unusual colors has stimulated research to determine the genetics underlying carrot color. Genes for carotenoid accumulation described by Gabelman’s group account for yellow and red color classes (Buishand *et al*., 1979). Their efforts described seven major genes accounting for difference among orange, white, yellow, and red root color. More recently the *Y* and *Y2* genes were mapped, a SCAR marker developed for *Y2* (Bradeen and Simon, 1998), and 20 QTL mapped for carotenoid content (Santos and Simon, 2002). A single major gene, *P1*, confers purple storage root color but this gene only accounts for part of the variation observed for purple color, as a wide range of pigmentation patterns occur, and at least one other gene, *P2*, influences pigmentation in aerial plant parts (Simon, 1996). To develop breeding stock with potential commercial application, carrot breeders utilize traditional regional carrots and long-ignored heirloom cultivars with unusual colors in crosses with adapted, good-flavored orange carrots to combine unusual color with acceptable flavor for modern consumers (Erdman *et al*., 2020).

Nitrates are important for their anti-nutritional value, especially for carrots used to make baby food. The inheritance of nitrate content in carrot is complex with incomplete dominance so that low-nitrate parents are necessary to obtain low–nitrate hybrids. In fact, while heterosis has significant positive effects upon many carrot production attributes, it is not observed for carotenoid or nutrient content, as mid parent values are observed in the majority of hybrids (Philipp *et al*., 2020).

Carrot flavor is a very important variable influencing consumer decisions. Flavor differences were noted between purple and yellow carrots hundreds of years ago and among modern orange carrot root types today, sweet and juicy flavor can be found in a wide range of types such as ‘Nantes’, ‘Kuroda’ and ‘Imperator’. With a broad genetic range in carrot flavor and the development of high value carrot products, including lightly processed “baby” or “cut and peel” carrots, improved raw carrot flavor has become a major breeding goal of carrot breeders in North America (Simon, 2000). Sweet flavor and succulent juicy texture are two of the major targets for improving raw carrot flavor. In addition to these two attributes, lack of harsh or turpentine flavor, caused by volatile terpenoids is the primary flavor component evaluated in selecting for improved flavor since high levels in harsh carrots can mask sweet flavor. Laboratory –facilitated selection is sometimes used for sweetness, using refractive index, colorimetric, or HPLC methods to quantify sugars; and for harsh flavor, using gas chromatography to quantify volatile terpenoids (Simon *et al.,* 1982).

The genetics of raw carrot sweet and harsh flavor has been described and the patterns of inheritance are complex. Sweet flavor, not surprisingly, is associated with higher sugar content which is polygenic, although a single major gene, *Rs*, determines whether reducing sugars glucose and fructose, or sucrose, are the primary storage carbohydrates (Stommel and Simon, 1989). While texture is an important component of raw carrot flavor, little attention has been paid to the genetics of this trait. Since variation in texture interacts with perception of sweetness and harshness, breeder selection of carrot flavor generally relies upon tasting roots in the field and/or during the period they are being stored for verbalization. Relatively little change occurs in carrot flavor or carotene content during early post-harvest storage so it is a convenient time to evaluate quality attributes. Unfortunately, the brittleness that accompanies crisp texture tends to have a negative impact on the “durability” of carrots in mechanical harvesting and washing (Philipp *et al.,* 2020).

#### 2.2 Nutritional Value of Carrot

##### 2.2.1 Bioavalaibility of β-Carotene

Deficiency in Vitamin A remains a major nutritional problem in most economically disadvantaged areas of the world (Olson 1994a, Sommer *et al.,* 1996), this makes the population to rely on dietary sources of provitamin. A carotenoid to meet the need of vitamin A. It has been considered that the most appropriate solution to this problem is the strategies developed by Public health which enhanced the increased intake of carotenoid rich vegetables and fruits (Solomon and Bulux 1993). Various factors affect the bioavailability of carotenoids, such as characteristics of the food source, interaction with other dietary factors and various subject characteristics (Bowen *et al.,* 1993, Erdman *et al*., 1993, Olson 1994b, Parker 1996), Size of the particle, the location of the carotenoid in the plant source (i.e. the pigment protein complexes of cell chloroplasts vs. the crystalline form in chloroplasts). Factors that affect proper micelle formation are included in characteristic that can affect carotenoid uptake and absorption (Erdman *et al*., 1993, Rock *et al* 1992, Zhou *et al.,* 1996). However, suggestions have been made that heat treatment may improve the bioavailability of carotenoids from vegetables (Poor *et al.,* 1993). During feeding of processed vs. raw vegetables the percentage changes in plasma of cis-β-carotene and α-carotene concentration remains the same. Daily consumption of processed carrots within 4 weeks will result production of plasma β-carotene response compare to the consumption of the same amount of the raw vegetables. Study has shown that thermal processing of this vegetable had substantially increased the proportion of cis-β-carotene isomers. Result from studies have also made a suggestion that isomers of cis-β-carotene have less of provitamine A activity than that of all-trans-β-carotene, and lower bioavailability may also be explained by some absorption and discrimination of isomers (Erdman *et al*., 1993, Gaziano *et al.,* 1995, de Pee *et al*., 1995). Consumption of food riches in carotenoid that have been treated with mild heat has sometimes but not always have been observed to enhanced the serum β-carotene or retinol concentration in population whose marginal vitamin A status is poor than (Bulux *et al.,* 1994, de Pee *et al.,* 1995, Solomon *et al.,* 1993, Solomon 1996). The following are factors that can seriously affect carotene absorption: high rates of parasitic infections, very low-fat diets consumption, and impaired absorption capability as a result of malnutrition (Bowen *et al.,* 1993, Erdman *et al*. 1993, Olson 1994b, Parker 1996).

##### 2.2.2 Calcium Transport Activity in Carrot

Intake of low dietary calcium can impact health negatively and enhanced the risk of diseases known as osteoporosis. Fruits and vegetables offer a diverse mixture of nutrients that promote good health, and it is generally believed that they will be more beneficial to human health than dietary supplements. One way to increase the nutrient content of some vegetables is to increase their bioavailable calcium levels. Carrots are among the most popular vegetables in the United States and contain high levels of beta carotene (the precursor to Vitamin A) and other vitamins and minerals; however, like many vegetables, they are a poor source of dietary calcium. By engineering carrots and other vegetables to contain increased calcium levels, one may boost calcium uptake and reduce the incidence of calcium deficiencies (Roger *et al*., 2007).

Generally, calcium (Ca) levels in plants can be engineered through high-level expression of a deregulated Arabidopsis calcium transporter. An Arabidopsis vacuolar calcium anti porter, termed Cation exchanger 1 (CAX1), contains an N-terminal auto inhibitory domain. Expression of N-terminal truncations of CAX1 (sCAX1) in plants such as potatoes, tomatoes, and carrots increase the calcium content in the edible portion of these foods. Presumably, these sCAX1 -expressing plants have heightened sequestration of calcium into the large central plant vacuoles. (Roger *et al.,* 2007. Modification of carrots to express increased levels of a plant calcium transporter (sCAX1), and these plants contain higher calcium content in the edible portions of the carrots, helps to improve the bioavailable calcium content of a staple food; when applied to a wide variety of fruits and vegetables, this strategy could lead to more calcium consumption in the diet. By this means one could rid of low intake of calcium in a deficient population. (Roger *et al.,* 2007)

##### 2.2.3 Storage and Preservation of Carrots

Garden vegetables lose their physiochemical and organoleptic properties in a few days after harvesting especially when they are stored in ambient conditions (Caron *et al.,* 2003). In carrots, mass loss and the incidence of disease in the root are the principal causes of postharvest loss during storage and commercialization (Oliveira *et al.,* 2001). In most vegetables, mass losses of 5% or higher can produce wrinkling and a consequent decline in consumer acceptance. This is due to high rates of transpiration, which affects the product’s appearance by wrinkling and altering the texture of its skin, among other effects (Caron *et al.,* 2003). The water content of carrot roots varies from 85 to 90%, a large part of which is lost through transpiration. Transpiration is a consequence of vapor pressure deficit (VPD), which results from the difference between the humidity at the surface of the product and the humidity of the surrounding air (Chitarra, 2005). Devraj (2001) emphasizes that 25-30% of the production of fruits and vegetables are wasted due to the lack of proper postharvest handling and storage. Carrot is well-storable vegetable species (Valšíková *et al*., 2009). The shelf life of carrot quality is ranged from 3 to 6 month at the temperature from - 0,5°C to +1,5°C (Valšíková *et al*., 2002), Uher *et al*., 2009) indicate that carrot designed for storage requires high relative air humidity because its anatomical structure does not allow preventing to water losses effectively. Carrot should be stored at relatively humidity of 98-99%. The useful life of product, e. g. carrot can be extended by using flexible plastic film that acts as modified atmosphere packaging. The aim of plastic film is to reduce the respiration, defend to the weight loss and microbial growth rates, as well as delay enzymatic deterioration, with the end effect of prolonging shelf life (Kumar *et al*., 1999), (Caron *et al*.,2003) also stated that package is very important factor affecting to the weight loss and storage period of carrot roots. (Oliveira 2001) found that the most suitable package material, from aspect of weight loss, is PVC film. On the other side, (Ayub *et al*., 2010) observed a higher percentage of carrot roots sprouting when stored wrapped in PVC film. (Koraddiand *et al.,* 2011) examined the effect of various types of packing materials with several vegetable species in refrigerator. They also confirmed the important role of package from aspect of weight loss and shelf life of stored products (Philipp *et al.,* 2020).

#### 2.3 Bacteria

Bacteria are most easily studied in pure cultures in which only a single species is present. Pure cultures were originally produced by limiting dilution in liquid medium. Today pure cultures are usually prepared on medium solidified with agar, a gelling agent derived from seaweed. A mixed bacterial suspension is mechanically spread on the agar surface to yield isolated individual bacterial cells. These grow to yield macroscopic colonies (clones) that can be used to prepare pure cultures. The ability to prepare pure cultures led to the study of bacterial classification and taxonomy. The first basis for classification was shape. Round bacteria are called cocci (singular coccus). Rod shaped bacteria are called bacilli (singular bacillus). Other shapes will be considered later in the course. Bacteria are very difficult to study microscopically unless stained. The staining characteristics of bacteria in the Gram stain are very useful in classification. Gram positives are violet, while gram negatives are red. Bacterial taxonomy today depends upon the extent of DNA sequence homology. An important laboratory technique for the amplification and detection of specific DNA sequences (as, for example, in a bacterium or a virus) is the polymerase chain reaction (PCR). Examples of when PCR is used for clinical diagnostics will be considered later in this course. However, for routine laboratory diagnosis the most important bacterial characteristics are:

i. The morphology of colonies on appropriate agar medium.
ii. Microscopic morphology and staining of individual bacteria.
iii. Simple biochemical characteristics such as the ability to ferment a given carbohydrate.
iv. Specific antigens detected by known antisera.

##### 2.3.1 Pathogenicity of Bacteria

Pathogenic bacteria utilize a number of mechanisms to cause disease in human hosts. Bacterial pathogens express a wide range of molecules that bind host cell targets to facilitate a variety of different host responses. The molecular strategies used by bacteria to interact with the host can be unique to specific pathogens or conserved across several different species. A key to fighting bacterial disease is the identification and characterization of all these different strategies. The availability of complete genome sequences for several bacterial pathogens coupled with bioinformatics will lead to significant advances toward this goal (Wilson *et al*., 2002).

Recently, significant evidence has emerged which indicates that markedly different microbial pathogens use common strategies to cause infection and disease. For example, many diverse bacterial pathogens share common mechanisms in terms of their abilities to adhere, Invade, and cause damage to host cells and tissues, as well as to survive host defenses and establish infection. Many of these commonalities of infection appear to be related to the acquisition of large blocks of virulence genes from a common microbial ancestor, which can be disseminated to other bacteria via horizontal transfer. A more thorough comprehension of the common themes in microbial pathogenicity is essential to understanding the molecular mechanisms of microbial virulence and to the development of novel vaccines and other therapeutic agents for the treatment and prevention of infectious diseases. The following are the virulence factor of bacteria: capsule production, presence of cell wall, production of toxins, adhesive ability, intracellular lifestyle, ability to invade the host (Wilson *et al*., 2002).

##### 2.3.2 Meaning of Isolation and Identification

In microbiology, the term isolation refers to the separation of a strain from a natural, mixed population of living microbes, as present in the environment, for example in water or soil flora or from living beings with skin flora, oral flora or gut flora, in order to identify the microbe(s) of interest. Historically, the laboratory techniques of isolation first developed in the field of bacteriology and parasitology (during the 19^th^ century), before those in virology during the 20^th^ century (Wikipedia 2021). Identification: Bacteria are classified and identified to distinguish among strains and to group them by criteria of interest to microbiologists and other scientists (Baron, 1996).

## CHAPTER THREE

### 3.0 MATERIAL AND METHODS

#### 3.1 MATERIALS

Conical flask, Measuring cylinder, Universal bottles, Test tubes, Spatula, Cotton wool, Foil paper, Petri dishes, Syringes, Glass slides, Jik, Test tube rack, Salmonella Shigella Agar (SSA), MacConkey Agar (Mac), Nutrient Agar (NA), Peptone water, Eosine Methylene Blue (EMB), Mueller Hinton, and Biochemical media.

#### 3.2 Media Preparation

Salmonella Shigella Agar (SSA), MacConkey Agar (Mac), Peptone water (PW), and Eosine Methylene Blue (EMB), were all prepared according to manufacturer’s instruction manuals and sterilized at 121°c for 15 minutes before it was used for the isolation of Pathogenic Bacteria from all the samples

#### 3.3 Collection of Sample

Twelve (12) samples of fresh carrots were purchased from different vendors at community market Abu Zaria. Samples were collected at two days’ interval and four samples per day were collected. The samples were placed in a new Polytene bags and then transported to the Department of Microbiology laboratory for analysis. All samples were processed within 24hours of collection.

#### 3.4 Isolation of Bacteria

##### 3.4.1 Serial Dilution

The epidermal of each of the carrots were scraped with the use of scalpel; 10grams of each of the samples were weighed into already prepared 90ml of peptone water (an enrichment medium) and were incubated for 3hrs. Serial dilution were made from the stock into five universal bottles containing 9mls of sterile distilled water each and labeled 10^1, 10^2, 10^3, 10^4, 10^5, with the use of sterile dropping pipette, 1ml of the sample in solution was pipetted into the first universal bottle, therefore making 10^1 dilution, this process was repeated serially to obtain 10^2, 10^3, 10^4, 10^5 dilutions (Public health England, 2019).

##### 3.4.2 Inoculating Procedure

Each of the samples, were pipetted using dropping pipetted from 10^4 dilution onto SSA, EMB, Mac, and a sterile bent glass rod was used to spread 0.1ml on it. After which it was incubated at 37°c for 24hr (Xianzhou *et al*., 2017).

##### 3.4.3 Microscopy

Growth was observed on each of the media after 24hrs of incubation based on morphology, texture, size, elevation, shape, and pigmentation of the colonies on the incubated medium.

###### 3.4.3.1 Procedure for Gram Staining

A clean grease free slide was taken. The smear of suspension was prepared on the clean slide with a loopful of sample. It was air dried and heat fixed Crystal Violet was poured and kept for 1 minutes and rinse with water. It was flooded with gram’s iodine for 1 minute and wash with water. Then, washed with 95% alcohol and rinsed with water immediately. Safranin was added for 1 minute and wash with water. Air dried, blot dried and Observed under Microscope. Discrete colonies of each of the samples were streaked onto a fresh nutrient Agar slant and incubated at 37°c for 24hr. The isolates were stored at 4°c for biochemical tests further identification (Philipp *et al.,* 2020).

#### 3.5 Identification of Bacteria Isolates

Biochemical test such as Indole test, Methylene red and Voges proskauer test, Citrate test (IMVIC). Triple sugar ion Agar, urease test and Catalase test were carried out for the identification as described below:

##### i. Procedure of Indole Test

Sterilized test tubes containing 4 ml of tryptophan broth were taken. The tubes were inoculated aseptically by taking the growth from 18 to 24hrs culture. The tubes were incubated at 37°C for 24hours. 0.5 ml of Kovac’s reagent was added to the 24hours broth culture. It was observed for the presence or absence of ring (Anupama, 2020).

##### ii. Procedure of Methyl Red (MR) Test

A clean test tubes containing 5ml of sterile MR-VP broth was inoculated with 24hrs culture It was incubated at 37°C for 24 hours. Following 24 hours of incubation, 0.5ml of methyl red indicator was added. It was observed for red color immediately (Sagar, 2018).

##### iii. Procedure for VP Test

The MRVP broth was inoculated with a pure culture of the isolate. It was incubated at 37°C for 24hrs. 6 drops of VP reagent I (alpha napthol) and 2 drops of VP reagent II (40% KOH) were added. It was observed for the color change (Crimson to ruby pink (red) color) in the broth medium (Sagar, 2018).

##### iv. Procedure of Citrate Utilization Test

Simmon citrate agar was prepared according to manufacturer’s instruction manual. A well-isolated colony is taken from 24hrs culture with a sterile inoculating wire loop. The citrate agar tubes were inoculated by streaking the surface of the slant. The cap of the test tubes was left loosened to ensure adequate aeration. The tubes were then incubated at 37°C for 24hrs. The test tubes were observed for change in color (Anupama, 2020).

##### v. Procedure of Catalase Test

A drop of 3% H2O2 was placed in the glass slide. A sterile needle was used to transfer a small amount of colony growth in the surface of a clean, dry glass slide. It was then observed for the evolution of oxygen bubbles (Sagar, 2018).

##### vi. Procedure for Urease Test

The surface of a sterile urea agar slant was streaked with a portion of 24hrs culture isolate. It was then incubated at 37°C for 24hrs. It was then observed for the development of a pink color (Sagar, 2018).

##### vii. Procedure for Triple Sugar Iron Test

With a straight inoculation needle, the top of isolate was picked, TSI was stabbed through the center to the bottom of the tube and then the surface of the agar slant was streak, it was then incubated at 37°C 24 hours. Then observed for reaction (Sagar, 2018).

Antibiotic Sensitivity Pattern of the Isolate

#### 3.6 Susceptibility Test Procedure

Sterile Petri dishes with Muller Hinton Agar was prepared. A pinch of the isolates was picked using sterile wire loop and dipped into sterile normal saline; the turbidity was compared with 0.5Macfarland standard. A sterile cotton swap was dipped into the inoculum and gently streaks the entire surface of the medium until evenly distributed to have a confluent growth on the petri plate. The inoculums were allowed to dry for 5 minutes along with lid in place. The discs were applied apart using aseptic technique. It was then incubated at 35°C for 24hrs after allowing the disc to diffuse within for some times. The plates were examined for zones of inhibition (Barth *et al.,* 2009).

### 4.0 CHAPTER FOUR

#### 4.1 Results

Below are the analyses of the result obtained from the twelve (12) carrots samples bought from community market Ahmadu Bello University Zaria. The result showed that pathogenic bacteria were isolated from all the carrot samples. The result shows that out of 12 samples analyzed *Proteus spp* was isolated from 5 samples (41%), *Citrobacter spp* was isolated from 3 samples (25%), *Klebsiella spp* was isolated from 2 samples (17%) and *Salmonella spp* from 2 samples (17%) as shown in fig 1 below.

**Figure 1:**
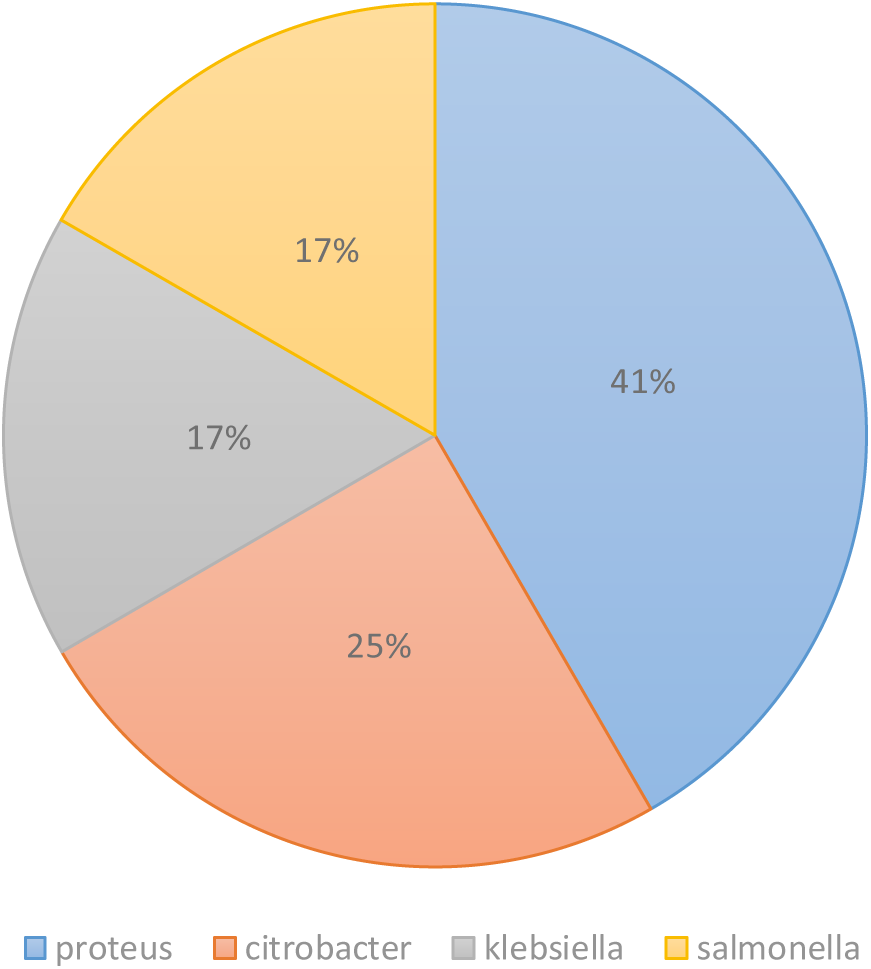
Occurrence of the Isolates in 12 Samples Examined from carrots sold in community market, Ahmadu Bello University, Zaria. The figure above shows the percentage occurrence of all the isolates in the samples examined, the highest percentage is shown in *Proteus* spp which was 41% followed by *Citrobacte*r spp which was 25%, *Salmonella* and *Klebsiella* which had the same percentage of 17%.

**Table 4.1:** Colonial morphology and Gram reaction of pathogenic bacteria isolated from carrot sold in community market Ahmadu Bello University, Zaria.

| Samples code | Shape | Elevation | Texture | Pigmentation | Gram Reaction |
| --- | --- | --- | --- | --- | --- |
| S1 | Circular | Raise | NM | Colorless | -ve rod |
| S2 | Circular | Raise | NM | Colorless | -ve rod |
| S3 | Circular | Raise | NM | Colorless | -ve rod |
| S4 | Circular | Raise | NM | Colorless | -ve rod |
| S5 | Circular | Raise | NM | Colorless | -ve rod |
| S6 | Circular | Raise | NM | Colorless | -ve rod |
| S7 | Circular | Raise | NM | Black Eye Dot | -ve rod |
| S8 | Circular | Raise | NM | Colorless | -ve rod |
| S9 | Circular | Raise | Mucoid | Pink | -ve rod |
| S10 | Circular | Raise | NM | Colorless | -ve rod |
| S11 | Circular | Raise | NM | Colorless | -ve rod |
| S12 | Circular | Raise | Mucoid | Pink | -ve rod |

**KEY: NM = Non-mucoid, -ve =Negative** The table above shows the colonial morphology and the gram reaction of the isolates. The colonial morphology was obtained based on the physical appearance of the colonies on the various plate used which are Eosin methylene blue agar, Salmonella shigella agar and MacConkey agar. The gram reaction was obtained by preparing a smear on a slide and stained then the stained slides were viewed under a microscope using 100X objective.

**Table 4. 2:**
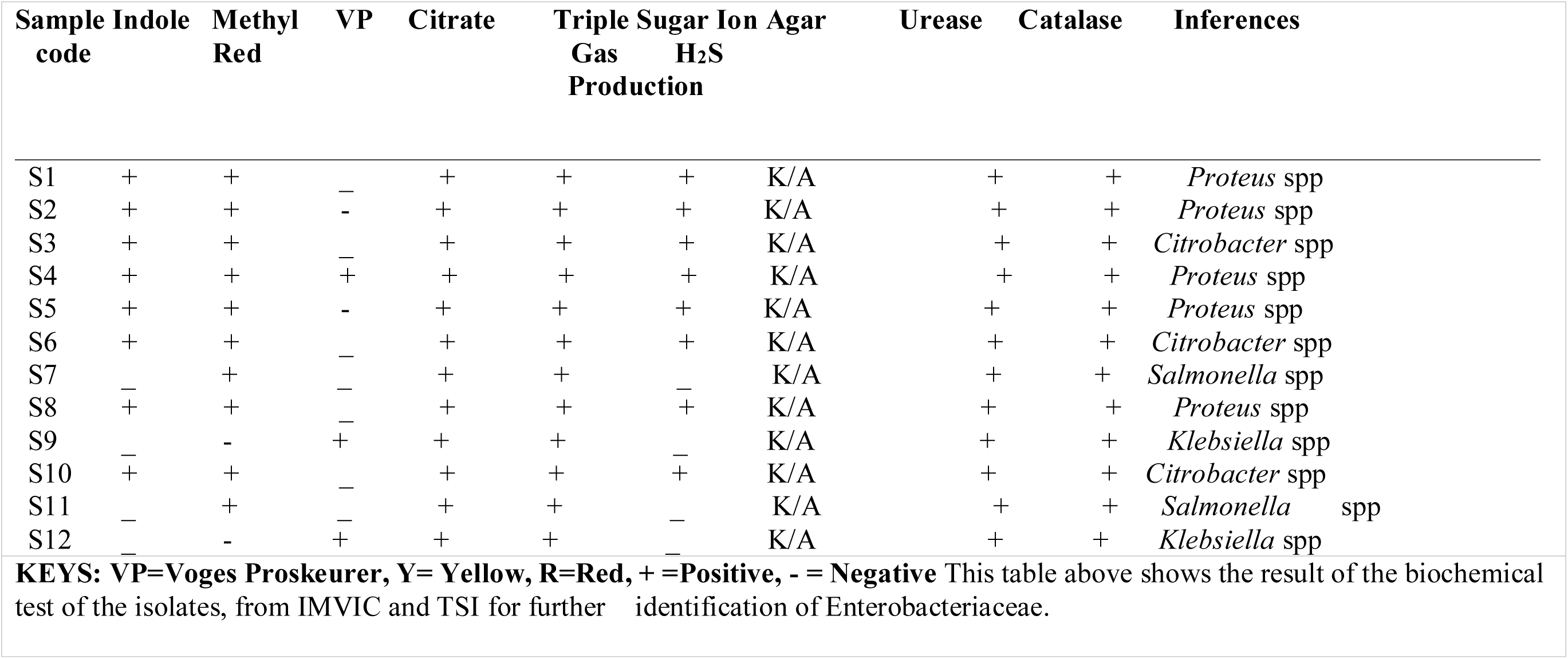
Biochemical characterization of pathogenic bacteria isolated and identified from carrot sold in community market, Ahmadu Bello University, Zaria.

**Table 4.2:** Antibiotic susceptibility pattern for the isolates from carrot sold in community market, Ahmadu Bello University, Zaria.

| **Antibiotics/Disc potency(µg)** | **Number of isolate tested** | **Number and percentage (%) of susceptibility** | **Number and percentage (%) of intermediate** | **Number and percentage (%) of resistance** |
| --- | --- | --- | --- | --- |
| Augmentin (10) | 12 | 6(50) | 3(25) | 3(25) |
| Amoxicillin (30) | 12 | 5(41.67) | 5(41.67) | 2(16.67) |
| Chloramphenicol (30) | 12 | 5(41.67) | 5(41.67)) | 2(16.67) |
| Ciprofloxacin (30) | 12 | 12(100) | 0(0) | 0(0) |
| Gentamicin (30) | 12 | 0(0) | 0(0) | 12(100) |
| Septrin (30) | 12 | 6(50) | 3(25) | 3(25) |
| Sparfloxacin (30) | 12 | 6(50) | 3(25) | 3(25) |
| Pefloxacin (30) | 12 | 6(50) | 3(25) | 3(25) |
| Tarivid (10) | 12 | 6(50) | 3(25) | 3(25) |

This table shows the antibiotic susceptibility pattern of the isolates which was calculated using the value gotten from measuring the zone of inhibition observed on Muller Hinton agar and it was calculated in percentage to determine the susceptibility, intermediate and resistance of the isolates to the antibiotics in line with CLSI 2020 as guide.

## CHAPTER FIVE

### 5.0 DISCUSSION CONCLUSION AND RECOMMENDATIONS

#### 5.1 Discussion

The research carried out revealed the following Bacterial (*Proteus* species*, Citrobacter* species*, Klebsiella* species, and *Salmonella* species) contamination of the carrot’s samples analyzed in various degrees. The percentage occurrence of the four organisms isolated from the carrot’s samples revealed that *Proteus species* had the highest occurrence of 41% in the carrot’s samples, followed by *Citrobacter spp* had 25%, *Klebsiella spp* had 17% and *Salmonella spp* had 17%.

The occurrence of *Protues spp* in the samples could indicate fecal contamination and poor hygienic environments. Proteus is the genus name belonging to the family of Enterobacteriaceae. It is an opportunistic human pathogen, found in human and gastrointestinal tract, skin and oral mucosa as well as feces, soil, water and plants (Yong *et al*., 2014). The effect of the organism could be food borne outbreak and urinary tract infection (UTI). *Proteus* spp (*proteus mirabilis and proteus vulgaris*) causes UTI and *proteus* UTI may give rise to bacteremia that are often fatal. Species of *proteus* has been suggested as a possible causative agent of outbreak of gastroenteritis resulting from the consumption of the contaminated food. Its role as the causative agent of acute gastroenteritis is very difficult to access, because of the high carrier rate in healthy individuals. (Samuel *et a*l.,2016). High level of susceptibility was observed in Ciprofloxacin followed by Augmentin and septrin and resistant to gentamicin

This study also showed that the sample was contaminated with *Salmonella spp* a food borne pathogen. The incidence rate of *Salmonella spp* in the samples analyzed was low. Consumption of vegetables contaminated with *Salmonella spp* could lead to light unrecognized microbial food borne hazards (Josiah 2015). Estimate has shown that each year two millions of people die of diarrheal diseases, mostly attribute to contaminated food and drinking water. Fresh produce is also susceptible to contamination during growth especially near sanitary facilities, harvest, and distribution. Salmonella contamination on the sample could also result from the market place where there are sold, there are usually kept on the ground and tables where flies uncontrollably perched on them before packaging for sales High level of susceptibility was observed in Ciprofloxacin followed by Augmentin and Septrin antibiotics.

The percentage occurrence of *Klebsiella* spp in the samples analyzed was 17% and it is a fecal coliform. *Klebsiella spp* is a rod-shaped non-motile, Gram-negative, lactose fermenting and facultative anaerobic bacteria which are usually found among the normal flora of skin, mouth, and intestines. Contaminated faeces during rain is washed into water bodies and even farmland where this vegetable is planted, the usage of this water without proper treatment on carrot processing will lead to it contamination. More so the main sources for this contamination are as follows; animal waste fertilizers, contaminated irrigation water and post-harvest washing using contaminated water by the food vendor. Isolation of *Enterobacteriaceae* and other bacterial species from vegetables has been reported by several researchers (Sahilah *et al*.,2010; Tunung *et al*., 2011).

Usually, the upper layer of the soil (30 cm2 from the ground) contains 106 - 107 bacteria/g as some farmers use animal manure or fecal as a fertilizer to enrich soil which could also be a source of contamination. On the other hand, contamination may occur through systemic contamination starting from the cultivation site to storage and handling. *Klebsiella* is found to cause infections in the urinary and lower biliary tract (Lopes *et al*., 2005; Ryan, 2004). *Klebsiella* is an opportunistic pathogen that primarily attack immunocompromised individuals and hospitalized patients. Improper food handling practices, poor hygienic condition of places where vegetables were displayed, use of contaminated equipment and containers during transportation can contribute to contamination with *Klebsiella* spp. According to some researchers (Ponniah *et al*., 2010; Tunung *et al*., 2010; Usha *et al*.,2010; Yang *et al*., 2008), poor hygienic practices and improper handling are considered as major factors for the contamination of food at the hypermarkets level. The minimum temperature for the growth of the mentioned pathogens is between 8 to 10°C so handling of carrot at a temperature below this can also regulate the level of *klebsiella spp* contamination. Coliforms are associated with both soil and decaying vegetation. Most of the *Klebsiella* spp isolated from carrots sold on campus were susceptible to Augmentin, Ciprofloxacin and Septrin. Microbial contamination of fresh vegetables with *Klesiella spp* is an important food safety concern. Consumption of contaminated fresh vegetables can represent a potential risk to consumer health, particularly in immunocompromised individuals. As mentioned before, *Klebsiella* is considered an opportunistic pathogen, so healthy adults are not considered to be at high risk of developing infections and illness. Isolation of *Klebsiella spp* from carrot vegetables has been reported by researchers (Puspanadan *et al*., Sahilah *et al*., 2010; Tunung *et al*., 2011)

In this study, it is obvious that most of the carrots sample frequently eaten raw on campus were contaminated with *Citrobacter* species in contrast to a research carried out at Ile-ife, Nigeria on Isolation and molecular characterization of *citrobacter* species in fruits and vegetables sold for consumption in Ile-ife, Nigeria where contamination of carrots sample by *citrobacter spp* was zero (Busayo 2019). Occurrence of *Citrobacter* spp in carrots sample depends on several factors such as: contact with the soil and water that are contaminated. Fruits and vegetables cultivated and harvested on the surface or in the soil are commonly contaminated due to contact with soil, manure, irrigation water, waste, and animal discharges. The antibiotic susceptibility of *Citrobacter* spp isolated from carrots sold on campus was susceptible to Augmentin, Ciprofloxacin and Septrin.

The implication of this is that consumptions of unhygienic raw carrots could be a source of resistance and virulence genes for both humans and other flora. *Citrobacter* spp recovered from carrots in this study are not a flora to carrots in particular; they are frequently isolated from animals and as an opportunistic pathogen in humans. *Citrobacter* spp are frequently been recovered in the intestines of animals and humans (Rogers *et al.,* 2016). It is possible that carrot might have been contaminated indirectly via fecal bacteria from animals during fertilization processes or by direct contact with humans during harvesting, handling and packaging, since direct contact with these produces is inevitable.

#### 5.2 Conclusion

It was found out that *Proteus* species*, Salmonella* species*, Klebsiella* species and *Citrobacter* species were present on carrots sold in Community Market Ahmadu Bello University Zaria Main Campus in various degrees. Carrots are root vegetables that promote good health and harbor a wide range of microbial contaminants that are of great concern. This now means that consumption of carrots sold in these markets can constitute a serious risk to health. In addition to that, the antibiotics susceptibility test on the isolates shows high level of susceptibility in Ciprofloxacin (100%) and Septrin (50%) for all the isolates.

#### 5.3 Recommendations

I. In order to avoid food-borne disease risk, special attention must be paid to improvement and control of the hygienic quality of fresh carrots such as: Hand washing, epidermal scrapping, thorough washing should be practiced by both the seller and the consumer; these will reduce the microbial load on carrots to minimal. Finding *Klebsiella spp* in raw food purchased at these markets indicates that additional food safety measures need to be practiced downstream, at retail and in the home.
II. *Klebsiella* cannot be easily dislodged from the surface of vegetables by gentle washing. Washing vegetables using disinfectants is one of the suggested ways to reduce risk. In addition, the government should survey vegetables seller and caution them on quality hygiene in order to minimize the risk of disease outbreak associated to consumption of contaminated food.
III. Potassium permanganate is as antimicrobial compound that is used in washing process of vegetables and is used to indicate if the rinsing procedure is correctly accomplished.
IV. The buyer and the consumer should be educated on the various sources of microbial contamination of vegetables and the effect of using polluted water to wash vegetable or not washing at all before eating and the use of unclean packaging materials and the need for proper sanitation of the surroundings where vegetables are sold.

